# Condition manipulation reveals an increase in sex-specific additive genetic variance, and reduced intersex genetic covariances in *Drosophila prolongata*, a species with sexual trait exaggeration

**DOI:** 10.64898/2026.06.18.733205

**Authors:** Tyler Audet, Sanhavy Vadivel, Aaron Taylor, Domenique Ammendolia, Noor Daanish, Olivia Beghin, Richard Yang, Saimieraa Yogaraajah, Ian Dworkin

**Affiliations:** Department of Biology, McMaster University, Hamilton, Ontario, Canada

## Abstract

The between sex genetic correlation for traits has long been hypothesized as a constraint to the evolution of sexual dimorphism. Both empirical and theoretical work has suggested that this constraint is influenced by genotype-sex-environment interactions. We examine genotype-sex-environment interactions in both sexually exaggerated and non-exaggerated legs of *Drosophila prolongata,* to examine the role of organismal condition on evolvability of an extreme trait. We employed a nested full-sib half-sib crossing design, providing food either *ad libitum*, or restricting food during larval growth, to each brood. When provided food *ad libitum*, inter-sex genetic correlations between traits is high and positive, whereas under food restriction this correlation substantially weakens, with a modest negative sign. Similarly, comparisons of the **G** matrix across sexes becomes less associated under food restriction. We discuss these results in the context of the growing appreciation of the factors that facilitate sex-specific evolutionary change.

## Introduction

The intersex genetic correlation (*r_MF_*) has been hypothesised to constrain one sex reaching a phenotypic optimum for a sexually dimorphic trait if there is countervailing selection on the opposite sex [1]. Directional selection on one sex can result in a concordant response in the other, supporting the idea that these correlations can influence the capacity for a system to evolve in a sex-specific manner [2,3]. The impact of consistently positive, and generally strong *r_mf_* (Poissant 2010), when each sex has distinct fitness optima, is this basis for the concept of intra-locus sexual conflict [1]. When only considering *r_mf_,* and if the additive genetic variance for the trait is similar in females and males, this will slow down rates of sex-specific adaptation, and is the basis for the so-called gender load [4]. These potentials impacts of a high, positive *r_mf_* can seem at odds with the common occurrence of subtle, and occasionally extravagant sexual dimorphism in nature [5], with some traits even being sex-limited [6,7]. Artificial selection on sexual dimorphism has demonstrated genetic variance for dimorphism using family based designs, often with a rapid evolutionary response [8,9]. In comparison, mass selection approaches, which are likely more comparable to transmission of alleles in natural populations, show the response to be slow for sexual dimorphism on body size, relative to sexually concordant selection for increasing or decreasing size [10].

It is important to determine the factors contributing to the relative degree of genetic constraint or independence on the evolution of sexual dimorphism for complex traits. There are two main genetic parameters influencing the rate of response to sex-specific selection, the intersex genetic covariances or correlations (such as *r_mf_*), and sex differences in the additive genetic variance for the trait. Even under a model where additive genetic variances for a trait is similar between the sexes, unless *r_fm_*=1, sex-specific responses occur, albeit more slowly when *r_mf_* is high. When quantified, *r_MF_* is generally positive [11], and the relationship between *r_MF_* and magnitude of sexual dimorphism is weakly negative [11,12]. This suggests that more sexually dimorphic traits are less constrained by r*_MF_*, at least after dimorphic phenotypes have evolved. Similar patterns are observed between sexual dimorphism and the intersex genetic correlation for induced mutations as well [13]. *r_MF_* itself may evolve, Delph et al. [8] selected for a reduction in *r_MF_* in the flower *Silene latifolia*, and found that low r*_MF_* treatments more readily responded to subsequent selection on reversal of sexual dimorphism.

The impact of the genetic correlation on reducing the rate of response to selection may be less severe when there are large sex-specific additive genetic variances for traits [1,14]. Although the genome is largely shared, sex-specific *V_A_* can arise through several routes, including changes in sex-specific regulatory mechanisms, dominance interactions, or environmental cues (recently reviewed in [15]). A meta-analysis looking at sexual dimorphism in *V_A_* for sexually dimorphic traits suggests that although, on average, *V_A_* is similar between the sexes, there are specific examples in which sexual dimorphism for *V_A_* can be higher, and that *V_A_* has a weak positive relationship with trait sexual dimorphism [16]. An important take away from these meta-analyses [11,16], is that quantitative genetic measures for complex traits are trait and population specific. Further, these genetic effects for complex traits, including dimorphism, depend on environment and organismal condition [15].

A univariate approach, such as examining *r_MF_* for a single trait, as a means of understanding genetic constraints on sexually dimorphic evolution does not reflect cross-trait, cross-sex covariances. Multivariate approaches suggest that constraints imposed by *r_MF_* may be less of an issue as (co)variance within and between traits within and between the sexes can provide variation for selection to act on. The inclusion of (co)variance matrices within sex (**G**_M_ and **G**_F_) and between sexes (**B**_MF_) can shift female response to selection to be more concordant and less of a constraint in response to selection ([17], as an example in *Drosophila serrata*). Within each sex, PC1 of **G** (**g**_max_) is the direction of greatest evolvability within that sex [18]. This direction of highest potential response is mediated by the covariance within **G**. When looking at gene expression, *r_MF_* and the off-diagonal (covariances) of **G** appear to be consistent predictors of dimorphism [19]. The extent to which sexual dimorphism can evolve is dependent on the similarity between **G** and **B** [1]. Specifically, how these matrices evolve to potentially facilitate sexually dimorphic evolution is still an open question. Sex specific **G** plays an important role in the response to antagonistic and concordant selection, and the evolution of sexual dimorphism [19]. Cross-trait asymmetries in **G** promote sexual dimorphism as an indirect response to concordant selection, while symmetries in **B** influences the response to antagonistic selection [19]. The intersex covariance within traits on the diagonal of ***B****_MF_* account for 86% of the predicted effect of concordant selection, and 95% of the predicted effect of discordant selection [20]. Thus, it has been suggested that indirect response to concordant selection due to covariances in **B** may initiate the evolution of sexual dimorphism from a monomorphic state [21]. As an example, in *D. melanogaster* Ingleby et al. [22] found that there was far less variance in **B** for cuticular hydrocarbons that were highly dimorphic, relative to CHCs that were less dimorphic.

Genetic variance in traits is also influenced by environment and condition. In particular, exaggerated secondary sexual traits tend to show increased condition dependent phenotypes, meaning that under lower condition or increased stress, these traits tend to decrease in size at a greater rate than non-sexually dimorphic traits [23,24]. Traits in distinct environments can be influenced by varying set of genes, making them analogous to different characters when under distinct environmental conditions, leading to environment specific genetic (co)variances [25,26]. Wood and Brodie [27] conducted a literature review of **G** between conspecific populations as well as between novel environments and found that the environmental influence on the shape and direction of **G** was comparable to differences between populations. Holman and Jacomb [28] looked at **G** and its components, and as a testament of the challenging of estimating these parameters well, often had wide confidence intervals making inference difficult. However, they found that the intersex inter-trait covariances (off diagonal of **B**) may not impose constraint on the direction of evolution, but when animals were food limited, the within sex covariance (off diagonal of **G**_M_ and **G**_F_) did impose constraint but also increased **g**_max_ by 4.6x. In contrast, In *D. serrata*, environment (diet) seems to not have a large impact on genetic covariance for fitness [29]. However, also in *D. serrata*, Delcourt et al. [30] suggested that within sex genetic correlation for fitness was positive, and between sex genetic correlation was negative in both diets tested. In a non-*Drosophila* example, Simons & Roff [31] measured genetic correlations using a split-family design in the cricket *Gryllus pennsylvanicus* where F_0_ adults were mated and half of their offspring were reared in controlled growth chambers, while the other half were reared in field conditions. They found reduced genetic correlations for morphological and life history traits within the field condition compared to the lab, but high between sex genetic correlations in both the laboratory and the field. Suggesting that *r_MF_* was high even in naturally variable environments in this species.

*D. prolongata*, which shared a common ancestor with *D. melanogaster* ∼35 million years ago [32], is a useful model for exploring the multivariate genetic (co)variance structure of exaggerated traits. The prothoracic legs of *D. prolongata* are much larger in males than in females or in any Drosophila species to date [33]. These enlarged legs are used for both intra-sexual fighting, as well as inter-sexual signalling and mating behaviours [32,34]. Interestingly, despite this extreme, male-biased dimorphism, previous work [35] suggests that the most sexually dimorphic traits are not the most condition-dependent, as expected under the genic capture model [23]. Due to this extreme sexual dimorphism we predicted decreasing inter-sex genetic correlations for more sexually dimorphic traits, as has been previously predicted [36,37], and observed in meta-analyses [11,12,14]. The uncommon pattern of condition dependence in *D. prolongata* and the ability to conduct quantitative genetic analysis introduces a unique opportunity to compare genetic (co)variances within and between sex for both exaggerated secondary sexual traits in comparison to developmentally similar non-exaggerated traits.

We use a nested full-sib half-sib crossing design to estimate additive genetic variances, *r_MF_*, **G**_M_ and **G**_F_, for the sexually exaggerated foreleg, and for the much less dimorphic midleg, along with thorax length, in an outbred population of *D. prolongata*. For each brood we manipulated access to food during growth, to identify how changes in condition mediate changes in genetic (co)variances. Among high condition cohorts, we observe that individual *r_MF_* values are high, and sex-specific **G** are aligned. We also find similar additive genetic variance for all leg traits between the sexes. Among food restricted cohorts, however, we observe modest negative, and generally weak *r_MF_*, a general increase in sex-specific *V_A_* and a reduction in the alignment of **G_M_** and **G_F_**. We discuss our results in an evolutionary framework for the potential of dimorphic evolution in varying environmental conditions.

## Methods

### Animal husbandry and crossing

The *D. prolongata* population were founded from collections in the Sa Pa region of Vietnam (22°20’N, 103°52’E) in 2018 [38]. From this population, 200 females were collected and maintained at 19.5°C in multiple (but varying numbers) of bottles that were combined each generation to maintain diversity and with overlapping generations. The population was maintained for ∼3 years prior to the current study with the population always maintained over a census size of 200 individuals, typically closer to ∼1000 individuals. For ∼1 year prior to this experiment this population was maintained at 21°C on a 12:12 day night cycle with non-overlapping generations.

Previous work has established this species as amenable to genetic crosses. Perdigón Ferreira et al. [39] used a population derived from isofemale lines to test for the relationship between sexual dimorphism and condition dependence, and found this species to be an exception to previous hypotheses on that relationship. This previous work, however, did not examine additive genetic variance or between sex (co)variances of this dimorphism and condition relationship, so we conducted a full-sib half-sib nested cross using 60 sires each allowed to mate with three (different) virgin dams in plastic vials on a sucrose/yeast media (Recipe in [39]) over the course of three days. After dams were allowed to mate, sires were discarded, and each dam was placed into their own vial for egg laying. Dams were transferred each day into a fresh vial for six consecutive days. The eggs laid on days one, four, five, and six were allowed to develop fully on the media while eggs laid on days two and three were subjected to food deprivation during growth. Condition manipulations via food deprivation were conducted as outlined in Stillwell et al. [40] but explained here briefly. Eggs laid on days two and three were allowed to hatch and consume media *ad libitum* for the first two days of growth. On the third day of development, these larvae were removed from food by adding ∼5ml of 40% sucrose solution and agitating gently on a nutator for 20 minutes. Floating larvae were removed using a paintbrush and placed in fresh vials with only a moist cotton ball, corresponding to their dam and sire combination. These larvae were allowed to finish developing in the absence of food (∼3 days without food). Offspring adults from the high condition and low condition treatments were collected and stored in 70% ethanol after emergence.

Foreleg (prothoracic) and midleg (mesothoracic) were dissected in 70% glycerol (in 1x PBS) with a small amount of phenol added as a bacteriostatic. Legs were imaged on a slide, and a lateral image of the thorax was captured using a Leica MZ75 microscope with a Leica IC90E camera (nominally 50X total magnification). While images were being collected, the microscope malfunctioned, causing the lens to stick within the body of the scope. Using a micrometer, we determined this deviation was consistent (28.3% difference in measurement), and we adjusted measurements accordingly during analyses. To correct, we took all individual images that were outside of the normally distributed trait sizes and multiplied their respective measurements by the observed micrometer difference taken with the broken dissecting scope, and with the scope after repair. When checked after correction, the traits in the combined data showed the expected normal distribution for size.

### Analysis and statistics

Linear measurements were converted from pixels to *µm* and log_2_ transformed prior to modelling. Modelling of (co)variance was conducted for sire and dam nested within sire using MCMCglmm v2.36 [41]. Highest posterior density (HPD) intervals were used to for uncertainty of estimates. Traits were modeled jointly as a multivariate response. For ease of downstream computation, we used a cell-means contrast coding, rather than the default treatment contrast coding in R. To reduce computational issues and speed, we partitioned our analysis into four multivariate mixed models. The first two models (examining partitions of the data based on high and low condition), and had fixed effects for sex, as well as random slopes for sex with respect to sire random effects, and dams (nested within sire) random effects. With this model specification, the multivariate mixed model was fit on the subset of the data for high condition and separately for low condition. We also fit models with condition as a fixed predictor (and random slopes as described in the previous model). In this case the data was subset with respect to each sex. As a “default” prior we used an inverse-Wishart distributions, with **S** = 0.6 × **I**_9_ and *v* = 10^−5^ for sire and dam level effects, as well as unique residuals for either sex or condition, depending on the model. 0.6 was used as a scalar multiplier for the 9×9 identity matrix as it was approximately 50% of the average of the total phenotypic variances across the traits, which was more sensible than a default of 1. To assess inferential robustness to choice of prior, we varied *v* across several orders of magnitude to confirm that the variance components did not change substantially. The results of the variance components were consistent until *v* > 10^−3^. This is as expected, as the effects of the more informative prior pulled covariances towards zero of the prior as *v* increased. We also confirmed the results (per trait), with each individual trait, confirming that the genetic (co)variances estimated from individual traits were similar to those estimated in the joint multivariate model (and that the prior was not having undue influence on estimates). For the results presented in the model MCMCglmm was run for 5 × 10^6^ iterations, with a burnin of 10000 and thinning to every 20^th^ iteration. The (co)variance matrix as well as highest posterior density intervals for our sire level (co)variances for each trait within and between sexes as well as within and between diet treatments were extracted for downstream analysis.

As we used log transformed values of our linear morphometric measures, estimated additive genetic (co)variances from the nested full-sib half-sib design are approximately equivalent to the coefficient of additive genetic (co)variances (as outlined in chapter 5 of ref. [42]), and along with the genetic correlations, are presented in the analysis. This is appropriate in this study as we want to focus on proportional changes to evolvabilities. Additionally, various forms of stress can substantially impact *V_E_* (thus *V_P_*), which can result in mis-interpreting inferences about evolvability using *h^2^*. These additive genetic variances were calculated as V_A_ = 4V_s_ (V_s_ represents sire variance) [43]. For comparison of the additive genetic variance covariance matrices, we used the Krzanowski shared space/ “correlation”, measuring the degree to which the first 4 eigenvectors (as we had 9 traits) of the two matrices being compared span a similar subspace[44,45]. We computed this with the MatrixCompare function in evolQG v0.3-4 [46]. We checked the results with additional methods such as Krzanowski sub-space comparison (in evolQG and MCMCglmm) and random skewers (evolQG) confirming consistent inferences across methods. We focus our comparisons *r_mf_*, **G_F_** and **G_M_** as sexual dimorphism in **G** for individuals traits tends to be more similar that **B** and thus may be a more conservative measure of potential constraint.

### Genomic Diversity methods

This study was conducted on a recently lab adapted population of *D. prolongata* assumed to be genetically diverse as it has been used for experimental evolution [47] and other genetic studies [39]. However, in a previous study in another lab, this population had seemingly modest genetic variance for traits [39]. The outlined methods for measuring genetic traits such as *r_MF_* and **G** are dependent on a genetically diverse populations, so we first confirmed that this population demonstrated segregating allelic variation. We extracted DNA from 25 adult males using a Qiagen DNeasy kit (DNeasy Qiagen kit, Cat # 69506). DNA was pooled and sequenced on an Illumina NovaSeq 6000 by Genome Quebec (Centre d’expertise et de services, Génome Québec) to a depth of 50x for our pool of 25 individuals.

Trimming was conducted with BBduk version 39.06 [48], and mapping was conducted with bwa-mem version 0.7 [49] to genome assembly ASM3634697v1. Optical duplicates were marked and removed using samtools version 1.23 [50]. Indel realignment was done using abra2 version 2.24 [51], and samtools was used to create an mpileup before SNP calling was conducted using poolSNP version 1 [52]. From the vcf outputted from poolSNP, repeatmasker version 4 [53] was used to mask repetitive regions. Grenedalf version 0.6.3 [54] was used to calculate Watterson’s π as a measure of diversity in 500bp windows across the genome.

## Results

### Genomic diversity in our lab population was moderate

The population has a genome wide average Watterson’s π was 0.0046, with variation as expected, within and between scaffolds (Figure S1, supplemental file 1), and diversity in 500bp windows is available in supplemental file 2.

### Food restriction reduced leg size in both sexes, but females showed a greater condition response

Food restriction reduced leg sizes in both sexes. We found that females are proportionally more condition dependent than males in all traits (Figure 1), which, for some traits, is counter to previous observations in the species from the same founding population [39]. The most condition dependent trait in males was the width of both forefemur and foretibia, which are the most sexually dimorphic traits, consistent with Perdigon Ferrera et al. [39].

**Figure 1:**
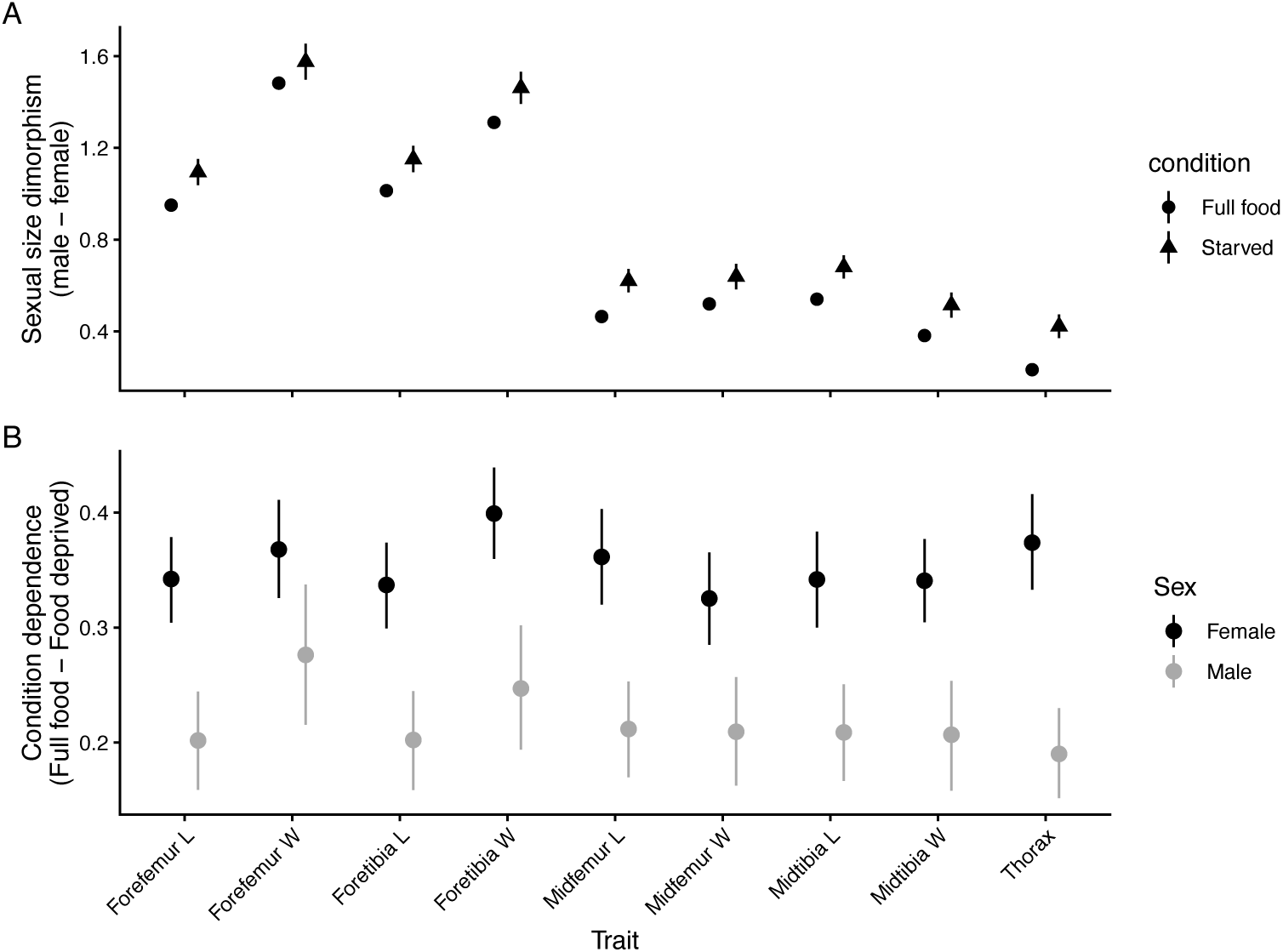
Length (L) and width (W) of all 4 leg traits (log_2_ transformed) measured on prothoracic, mesothoracic legs. A) male – female as a measure of sexual dimorphism, and B) full food – food deprived condition as a measure of condition response. Error bars represent 95% credible intervals from the posterior distribution (for all figures).

### Additive genetic variation increases for most traits under low condition, but there is no consistent pattern between degree of sexual dimorphism or condition dependence of traits, and amount of additive genetic variance

Additive genetic variance did not show a clear relationship with condition dependence in either sex, or in either condition (Figure 2A). When additive genetic variation was compared to trait sexual size dimorphism, our full food treatment did not show a clear relationship between SSD and V_A_ (Figure 2B). There was a potentially weak, although with high uncertainty, relationship between V_A_ and SSD in food deprived males (Figure 2B).

**Figure 2:**
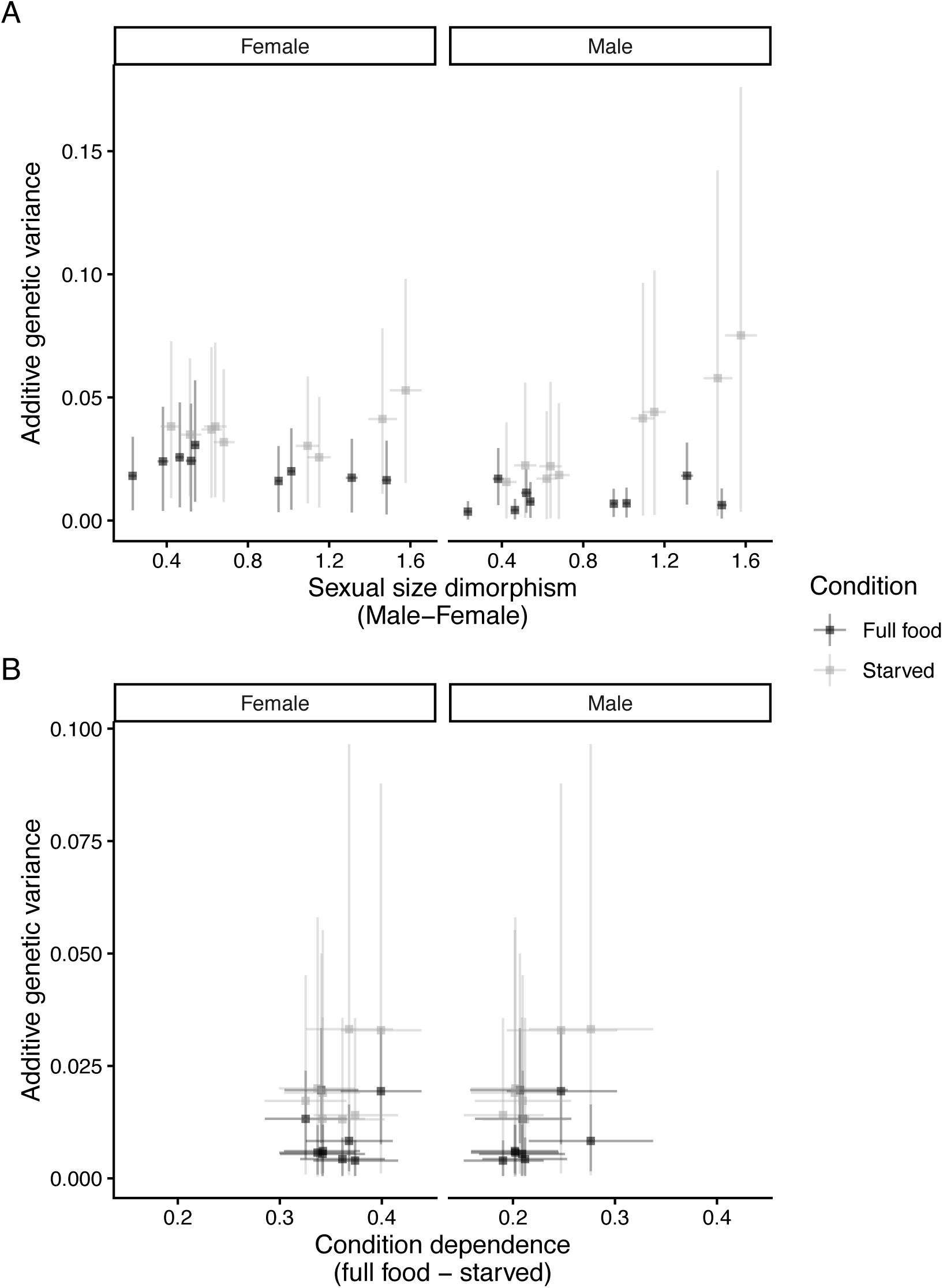
Association between additive genetic variance and sexual dimorphism (A) or condition response (B). For both sexes (split by panel) and each condition treatment. As data was modeled on log transformed values additive genetic variances approximate coefficient of additive genetic variances.

### r_MF_ and between sex Krzanowski correlation for the **G**-matrix is reduced in low condition cohorts

To examine changes in the potential for genetic constraint imposed by *r_MF_*, we measured *r_MF_* relative to the degree of SSD of each trait in both high and low condition (Figure 3). In the high condition treatment, *r_MF_* was positive and generally moderate to high for all traits. The relationship between *r_MF_* and the degree of trait SSD shows minimal association (Figure 3). In our low condition treatment, *r_MF_* was near zero for less sexually dimorphic traits (Figure 3). For the most sexually dimorphic traits (which correspond to foreleg traits), *r_MF_* is consistently negative (forefemur length: *r_MF_*= −0.5736, 95% CI=-0.981,0.0163; forefemur width: *r_MF_*= −0.6265, 95% CI=-0.989,-0.0715; foretibia length: *r_MF_*= −0.6554, 95% CI=-0.989,-0.1211; foretibia width: *r_MF_*=-0.5883 95% CI=-0.989, 0.0550). For both femur and tibia length, 95% confidence intervals do not overlap the high condition confidence intervals, while the width traits overlap between the two treatments, albeit very slightly for femur width which is the most sexually dimorphic trait (Figure 3).

**Figure 3:**
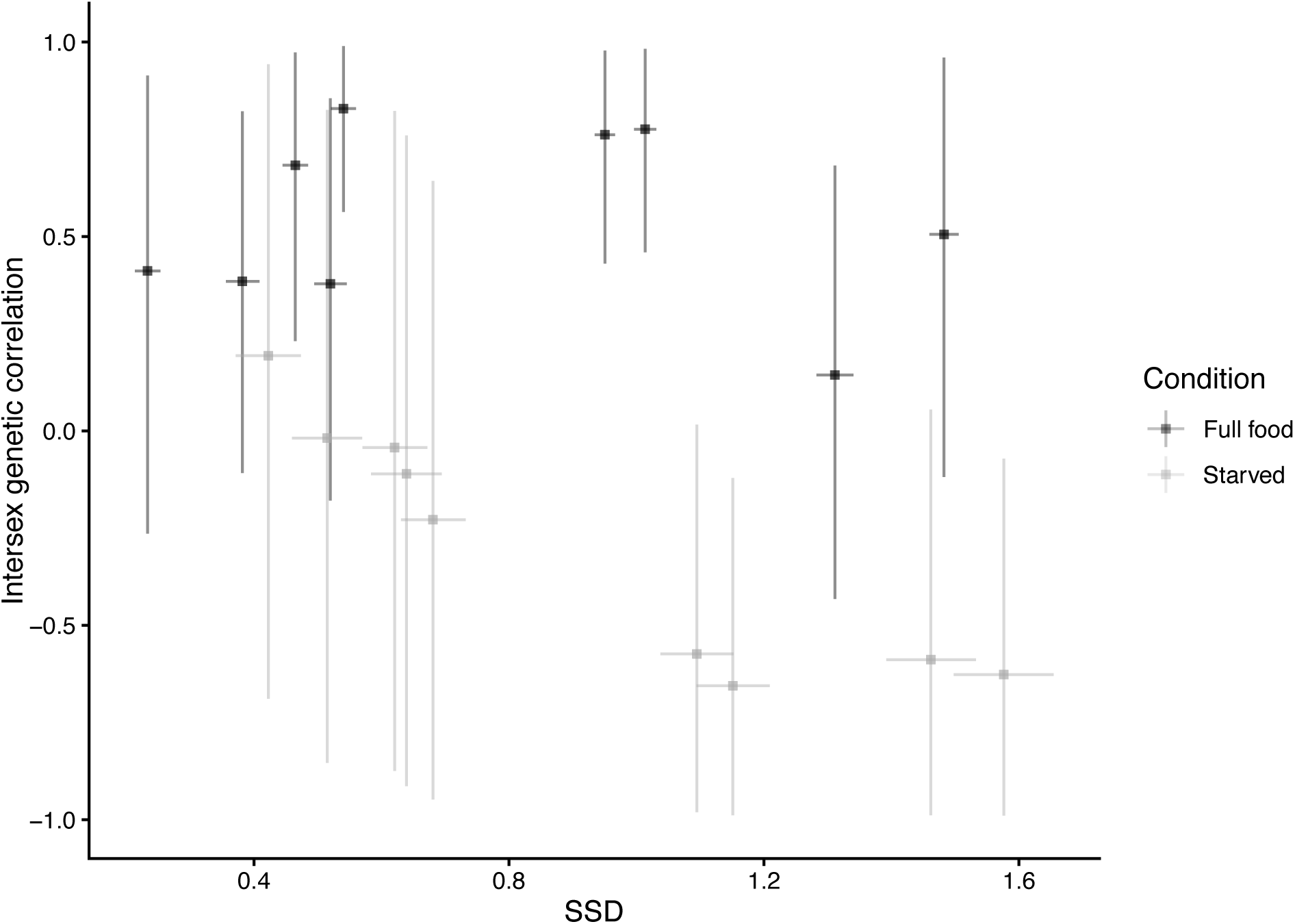
The intersex genetic correlation (*r_MF_*) and its relationship to sexual dimorphism (SSD) across traits, in both the full food and the food limited condition..

To examine this genetic constraint in a multivariate context, we looked at changes to the Krzanowski correlation between the conditions within sex and between the sexes within each condition. The correlation between **G**_F_ across high and low condition, and **G**_M_ in high and low condition was positive and similar (Figure 4; Females: 0.671 95% CI= 0.540,0.763; Males: 0.659 95% CI=0.528, 0.728). The **G**-matrix Krzanowski correlation between sexes in high condition was also positive (Figure 4; 0.711 95% CI=0.593,0.849). The **G**-matrix Krzanowski correlation between the sexes in low condition, although still positive, was lower than in high condition with non-overlapping 95% confidence intervals (Figure 4; 0.510 95% CI=0.398,0.591).

**Figure 4:**
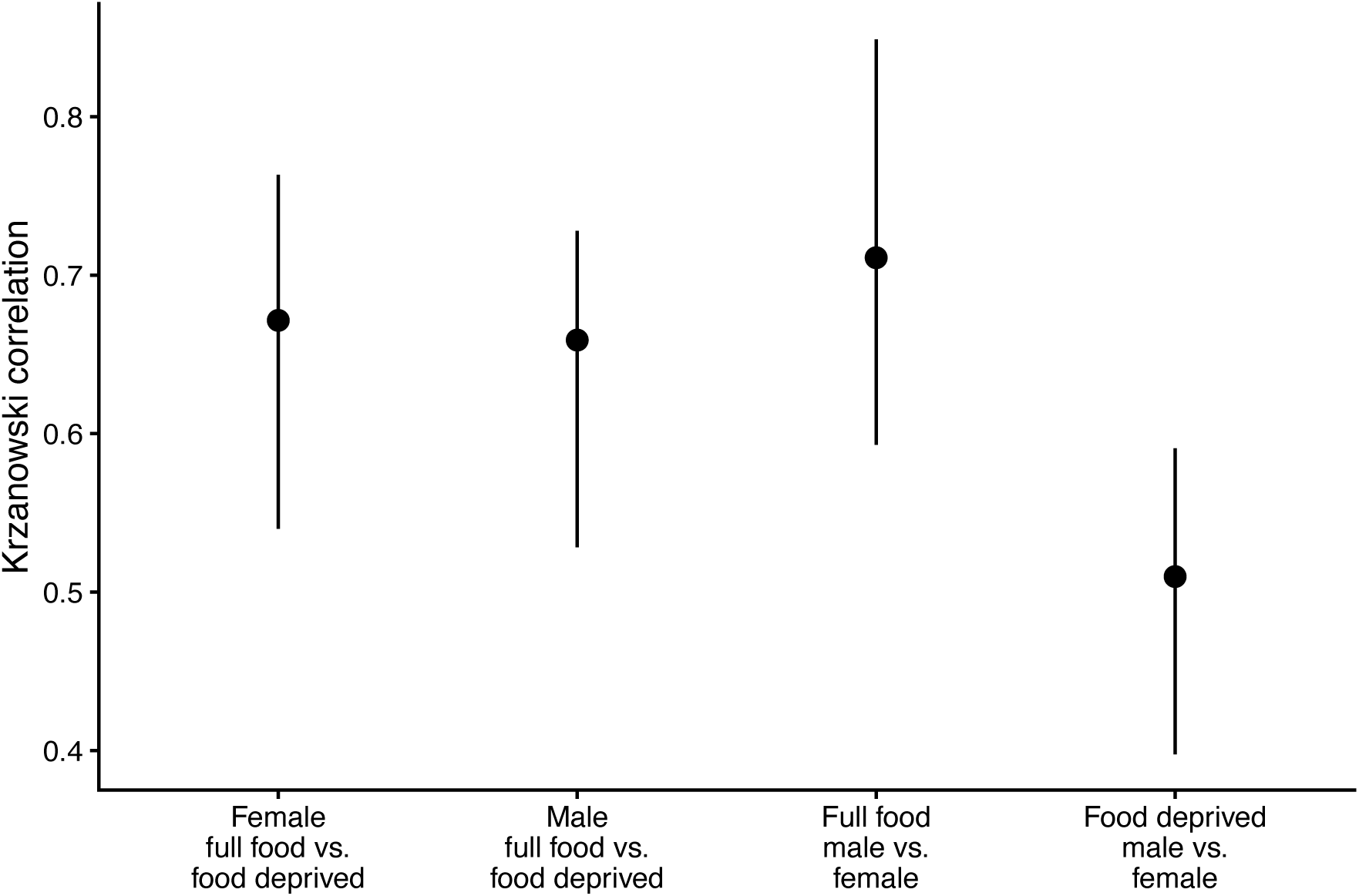
Krzanowski correlation between **G** matrices: **G**_F_ in full food vs. food deprived treatments, **G**_M_ in full food vs. food deprived treatments, **G**_M_ vs. female **G**_F_ in full food treatment, and male **G**_M_ vs. female **G**_F_ in food deprived treatment.

## Discussion

As the intersex genetic correlation for many traits (*r_MF_*) is quite high [11], it has been suggested it can be a constraint to the evolution of sexual dimorphism [1]. Yet, *r_MF_* is both evolvable[8], and context dependent [28,30,31]. Here, we demonstrate that for an exaggerated male secondary sexual trait, *r_MF_* not only changes under condition manipulation, but changes sign from positive in high condition to negative in low condition, specifically under reduced food availability (Figure 3). When *r_MF_* was experimentally altered in *Silene latifolia* by Delph et al. [8], they found that artificially selected lineages with reduced *r_MF_* were able to respond rapidly to sex specific selection compared to higher *r_MF_* lineages. In *D. prolongata*, response to selection on degree of sexual dimorphism may be constrained when conditions are consistently favorable and *r_MF_* is high for leg traits, but less constrained under more variable conditions. However, we still do not know whether changes to *r_MF_* occur before, during, or after sexually dimorphic evolution.

Another potential mechanism for sexually dimorphic evolution occurs via sexual dimorphism for additive genetic variance. Fairbairn et al. [14] found evidence of sex specific variance in autosomal genetic variance in the water strider *Aquarius remiges*, as well as trait- and sex-specific genetic dominance associated with sexual dimorphism. In *D. prolongata* however, additive genetic variance between the sexes, and in relation to the degree of sexual dimorphism were generally modest (Figure 2). If this pattern was present ancestrally, then differences in additive genetic variance were likely not a major contributor to the divergence in phenotype in this species.

We also demonstrate that **G** diverges between the sexes under condition manipulation. A reduction in *r_MF_* is not necessarily sufficient to allow dimorphic evolution, as (co)variance in **G**/**B** can also constrain or facilitate phenotypic divergence [55]. Our results again show that in high condition **G_M_** and **G_F_** are well aligned (Figure 4), however, under nutritional restriction, **G_M_** and **G_F_** show a reduced association suggesting a possible reduction in constraint.

Previous work looking at genetic correlations in *D. melanogaster* across an environmental cline, found consistently positive *r_MF_* for cold resistance and desiccation resistance which was inferred to increase adaptive potential in new environments [56]. This alignment of *r_MF_* under variable environments has been shown previously in laboratory environments as well in *Drosophila* [57,58]. While patterns **G** and **B** have been shown the change under environmental manipulation, to our knowledge this is the first demonstration of this involving a highly exaggerated sexually dimorphic trait.

Our results here add to our understanding of how shared genetic architecture of a trait between the sexes can produce variable effects depending on the growth environment. Framed another way, we would argue that recognizing the contribution of Genotype-Sex-Environment interactions along with the nature of variability in the environment itself may be key to understanding the rate at which sexual dimorphism evolves. However, many studies addressing these questions focus on traits that have already undergone substantial sexually dimorphic evolution, and find potentially inconsistent patterns for genetic constraint and its relationship to dimorphism (as a broad example [11]). We rarely know what *r_MF_*, **G**, or **B** looked like in ancestral populations at the initiation of divergence of phenotypes between the sexes, and how genetic constraints change after dimorphism has evolved. Future work looking at closely related species may help to infer to evolutionary history of this environment dependence genetic correlation. Here, we have shown that these parameters, often thought of as constraints, are more variable and environmentally dependent in a sexually exaggerated trait, and provide a potential route to more rapid evolution of sexual dimorphism.

## Author contributions

Experimental design: TA, ID

Experimental cross: TA

Dissection/imaging: SV, AT, DA, ND, OB, RY, SY

Image measuring: SV

Analysis: TA, SV, ID

Manuscript drafting: TA, SV, ID

## Acknowledgements and funding

This work was funded by National Science and Engineering Research Council Discovery grants (NSERC) to ID (2016-06453 & 2024-06914), and Ontario Graduate Scholarships (OGS) to TA.

